# Seroprevalence of Foot and Mouth Disease Virus Infection in some Wildlife and Cattle in Bauchi State, Nigeria

**DOI:** 10.1101/728501

**Authors:** Y.J Atuman, C.A Kudi, P.A Abdu, O.O Okubanjo, A Abubakar, H.G Ularamu, Y Wungak

## Abstract

**Background:** Foot and mouth disease (FMD) is one of the most economically important transboundary animal diseases with devastating consequence on livestock production and wildlife conservation. The objectives of the study were: to determine the seroprevalence of FMDV in wildlife and cattle and identify circulating FMDV serotypes in wildlife and identify potential risk factors that will contribute to transmission of the disease at the wildlife-livestock interface in Yankari Game Reserve and Sumu Wildlife Park in Bauchi State, Nigeria.

**Methods:** Blood samples were collected between 2013 to 2015 from some wildlife and cattle respectively within and around Yankari Game Reserve (YGR) and Sumu Wildlife Park (SWP) in Bauchi State, Nigeria. The Wild animals were immobilized for blood collection using a combination of Etorphine Hydrochloride (M99® Krüger-Med South Africa) at 0.5-2 mg/kg and Azaperone (Stresnil®, Janssen Pharmaceuticals (Pty.) Ltd., South Africa) at 0.1 mg/kg using a Dan-Inject® rifle (Dan-Inject APS, Sellerup Skovvej, Denmark) fitted with 3 ml dart syringe and for reversal, Naltrexone (Trexonil® Kruger-Med South Africa) at 1.5 mg IM was used, cattle were restrained by the owners for blood collection. Harvested Sera from blood were screened for presence of Antibodies against FMDV using prioCHECK® 3 ABC NSP ELISA kit and positive samples from wildlife were serotyped using Solid-Phase Competitive ELISA, (IZSLER Brescia-Italy). Data obtained were analysed using Graphpad Prism version 7.

**Results:** The results showed that 197 (65.7%) of the 300 serum samples from cattle and 13 (24.5%) of the 53 serum samples from wildlife tested positive for antibodies to the highly conserved non-structural 3-ABC protein of FMDV and statistically significant (P <0.05). Classification of cattle into breed and sex showed that detectable antibodies to FMDV were higher (P <0.05) in White Fulani 157 (72.8%) than red Bororo 23 (39.7%) and Sokoto Gudali 17 (33.3%) breeds of cattle whereas in females detectable FMDV antibodies were higher (P <0.05) 150 (72.8%) than in males 47 (50.0%). In the wildlife species, antibodies to FMDV were detected in waterbuck 2 (28.6%), elephant 1 (25.0%), wildebeest 4 (33.3%) and eland 6 (25.0%). Four serotypes of FMDV: O, A, SAT-1 and SAT-2 were detected from the 3-ABC positive reactors in waterbuck, elephant, wildebeest and eland. Contact of wildlife and cattle during utilization of the rich resources in the conservation areas is a potential risk factor for the spread of FMDV in the study area.

**Conclusions:** Presence of FMDV antibodies in cattle and some wildlife were observed and serotypes of FMDV: O, A, SAT-1 and SAT-2 were detected from the 3-ABC positive reactors in some of the wildlife. The study highlights the need for active surveillance of FMDV in wildlife and pastoral cattle within and around wildlife conservation areas in Nigeria. FMD surveillance system, control and prevention program that targets wildlife and livestock at the wildlife-livestock interface level will be beneficial to the livestock industry and wildlife conservation goals in Bauchi State, Nigeria.

**Author summary:** Foot and mouth disease (FMD) is an important trans-boundary viral disease of both domestic and wild cloven hoofed animals characterized by high morbidity with devastating consequence on the livestock worldwide. Despite the endemic nature of FMD in Nigeria, little is known about the epidemiology of the disease at the wildlife-livestock interface level. To address this gap, blood samples were collected between 2013 to 2015 from some wildlife and cattle respectively within and around Yankari Game Reserve (YGR) and Sumu Wildlife Park (SWP) in Bauchi State, Nigeria. Wild animals were immobilized using a combination of Etorphine Hydrochloride (M99® Krüger-Med South Africa) at 0.5-2 mg/kg and Azaperone (Stresnil®, Janssen Pharmaceuticals (Pty.) Ltd., South Africa) at 0.1 mg/kg using a Dan-Inject® rifle (Dan-Inject APS, Sellerup Skovvej, Denmark) fitted with 3 ml dart syringe and for reversal, Naltrexone (Trexonil® Kruger-Med South Africa) at 1.5 mg IM was used, cattle were restrained by the owners for blood collection. Harvested Sera from blood were screened for presence of Antibodies against FMDV using prioCHECK® 3 ABC NSP ELISA kit and positive samples were serotyped using Solid-Phase Competitive ELISA, (IZSLER Brescia-Italy). Out of the 300 and 53 sera collected from cattle and wildlife 197 (65.7%) and 13 (24.5%) (P <0.05) respectively tested positive for antibodies to the highly conserved non-structural 3-ABC protein of FMDV by the FMDV-NS blocking ELISA. Classification of cattle into breed and sex showed that detectable antibodies to FMDV were higher (P <0.05) in White Fulani 157 (72.8%) than red Bororo 23 (39.7%) and Sokoto Gudali 17 (33.3%) breeds of cattle whereas in females detectable FMDV antibodies were higher (P <0.05) 150 (72.8%) than in males 47 (50.0%). In the wildlife species, antibodies to FMDV were detected in waterbuck 2 (28.6%), elephant 1 (25.0%), wildebeest 4 (33.3%) and eland 6 (25.0%). Four serotypes of FMDV: O, A, SAT-1 and SAT-2 were detected from the 3-ABC positive reactors in waterbuck, elephant, wildebeest and elands. The results showed presence of antibodies to FMDV in some wildlife and cattle and suggest that wildlife could equally play an important role in the overall epidemiology of FMD in Nigeria. FMD surveillance system, control and prevention program should be intensified in the study area.

## Introduction

Foot and mouth disease (FMD) is one of the most economically important transboundary animal disease in the world caused by Foot and mouth disease virus (FMDV) a member of the genus *Aphthovirus* belonging to the *Picornaviridae* family (1). FMDV is a small non-enveloped virus and has a genome of 8.5 kb which encodes for structural proteins (VP1, VP2, VP3 and VP4) as well as non-structural proteins (NSPs) (2, 3).A structural protein produces antibodies to FMDV in vaccinated animals, whereas infected animals produce antibodies to both the structural and non-structural proteins (3) and assays to demonstrate antibodies against non-structural proteins have potential to differentiate infected from vaccinated animals (4,5,6,7). Seven immunologically different serotypes of the FMDV are known: O, A, C, Asia-1, South-African Territories (SAT) −1, −2 and −3, which comprise more than 65 subtypes (8).

The transmission of FMDV in sub-Saharan Africa is mainly driven by two epidemiological cycles: one in which wildlife plays a significant role in maintaining and spreading the disease to other susceptible wild and/or domestic ruminants (9,10). Whilst with the second cycle the virus is solely transmitted within domestic populations and hence is independent of wildlife (11). The disease is endemic in some parts of Europe, Africa, Middle East and Asia and has contributed to significant declines in wildlife and livestock populations in those regions (12, 13, 14, and 15). The first reported case of FMD outbreak in Nigeria was in 1924, which was attributed to type O virus (16). Subsequently, other serotypes (A, SAT 1 and SAT 2) were reported (17, 18, 19, 20, 21, 22) and recently SAT 3 serotype (23).

In spite of the annual FMD burden in Nigeria, sero-epidemiology and sero-typing studies for FMD infections are inadequate. The current trend of FMD occurrence in Nigeria showed that there are regular outbreaks, poor control measures and lack of enforcement of legislation guiding disease reporting to veterinary authority (24, 25). The presence of antibodies to FMDV in several wildlife species have been documented in studies conducted in different countries of Africa mainly eastern and southern regions (26, 27, 28). There has been limited monitoring of infectious diseases like FMD in wildlife in Nigeria. Domestic livestock sometimes do share the same range with wildlife in YGR and SWP in Bauchi State, Nigeria (29) and there is concern that wildlife may form a reservoir for FMDV. Consequently, there is need to understand the potential role of wildlife as reservoir of FMDV to aid in the design and implementation of the disease management programs. The aim of the study was to determine the seroprevalence of FMDV in wildlife and cattle and identify circulating FMDV serotypes in wildlife in YGR and SWP in Bauchi State, Nigeria.

## Materials and methods

### The study area

The study locations were Yankari Game Reserve (YGR) and Sumu Wildlife Park (SWP) in Bauchi State, Nigeria with human settlements surrounding them. YGR covers an area of about 2,244 square kilometres, it is an important refuge for over 50 species of mammals and over 350 species of birds and is one of the few remaining areas where wild animals are protected in their natural habitat in Nigeria (30, 31). SWP covers about 40 square kilometer area and habours species of wildlife including impala (*Aepyceros melampus*), springbok (*Antidorcas marsupialis*), oryx (*Orynx gazelle*), eland (*Taurotragus oryx*), zebra *(Equus quagga crawshayi*) kudu (*Tragelaphus strepsiceros*), blue wildebeest (*Connochaetes taurinus*), and giraffe (*Giraffa camelopardalis*) and is located about 60 km north of Bauchi the State capital (29).

### Serum sample collection

Field sampling was conducted between February 2013 to December 2015 and blood samples were collected from 300 cattle, and 53 wildlife including four elephant (*Loxodonta Africana*), eleven waterbuck (*Kobus ellipsiprymus*), one Hartbeest (*Alcelaphus buselaphus caama*) from YGR and twenty four eland (*Taurotragus oryx*), twelve blue wildebeest (*Connochaetes taurinus*) and one kudu (*Tragelaphus strepsiceros*) from SWP following chemical immobilization using Etorphine hydrochloride (M99® Krüger-Med South Africa) at 0.5-2 mg/kg and Azaperone (Stresnil®, Janssen Pharmaceuticals (Pty.) Ltd., South Africa) at 0.1mg/kg delivered intramuscularly (IM) using a Dan-Inject® rifle (Dan-Inject APS, Sellerup Skovvej, Denmark) fitted with 3ml dart syringe and barbed needles and for reversal Naltrexone (Trexonil® Kruger-Med South Africa) at 1.5mg IM was used. The serum samples were harvested from the blood into cryovials after spinning for 10 min at 1200 g and were divided into aliquots, labelled and kept at −20 °C until used.

### Detection of antibodies against FMDV non-structural proteins (NSPs) by ELISA

The ELISA was performed according to the manufacturer’s instructions (PRIOCHECK® FMD-3ABC NS protein ELISA) for detection of antibodies to the non-structural polypeptide 3 ABC of FMDV in serum which detects infected animals regardless of their vaccination status and the FMDV serotype that caused the infection (32). Briefly, 80 μl of the ELISA buffer and 20 μl of the test sera were added to the 3ABC-antigen coated test plates. Negative, weak positive and strong positive control sera were added to designated wells on each test plate, gently shaken and incubated overnight (18 h) at 22°C. The plates were then emptied and washed six times with 200 μl of wash solution and 100 μl of diluted conjugate was added to all wells. The test plates were sealed and incubated for one hour at 22ºC. The plates were then washed six times with 200 μl of wash solution and 100 μl of the chromogen (tetra-methyl benzidine) substrate was dispensed to all wells of the plates and incubated for 20 min at 22ºC following which 100 μl of stop solution was added to all the wells and mixed gently. Readings were taken on a spectrophotometer Multiskan® ELISA reader (Thermo Scientific, USA) at 450 nm and the OD 450 values of all samples was expressed as Percentage Inhibition (PI) relative to the OD 450 max using the following formula PI = 100 – [OD 450 test sample/OD450 max] × 100. Samples with PI = ≥ 50% were considered positive for FMD antibody while those with PI < 50% were declared negative for FMD antibody. Since the 3-ABC ELISA for FMD was = 100% specific and > 99% sensitive, the percentage prevalence was taken as true prevalence.

### Detection of FMDV specific antibodies using solid-phase competitive enzyme linked immunosorbent assay

The 3ABC ELISA positive serum samples were analyzed for FMD-specific antibodies using a Solid-Phase Competitive ELISA (SPCE) as previously described for serotypes O, A, SAT 1 and SAT 2 (32, 33). The assays were performed using antibodies FMDV ELISA kits for serotypes O, A, SAT 1 and SAT 2 produced by IZSLER Biotechnology Laboratory (Italy). Briefly, 96 wells pre-coated with FMDV antigens captured by FMD serotypes O, A, SAT 1 and SAT 2 specific MAb flat-bottomed plates were used. Four dilutions of sera at 1/10, 1/30, 1/90 and 1/270 were made. Without washing, the conjugate (Horse-radish peroxidase) was added and incubated at room temperature for 1 h. The plate was washed and the substrate/chromogen solution (tetra-methyl-benzidine) was added and kept in the dark for 20 min. The reaction was stopped by the addition of a stop solution and the plates were read on a MultiSkan® spectrophotometer ELISA plate reader (Thermo Scientific, USA) at 450 nm wavelength. Serum end-point titre was expressed as the highest dilution producing 50% inhibition, with serum having end point titre ≥ 50% being classified as positive for the specific FMD antibody.

Data obtained were analysed using Graphpad Prism version 7. Results were summarized in tables and expressed as percentages and levels of association between positivity and sex, breed, age and animal species were derived using Chi-square. Values of P ≤ 0.05 were regarded as statistically significantly different.

## Results

Overall seroprevalence of FMDV in wildlife was 24.53% (Table 1). Detectable antibodies to FMDV were observed in waterbuck 2 (28.6%), elephant 1 (25.00 %), wildebeest 4 (33.3%) and eland 6 (25.0 %).

**Table 1:**
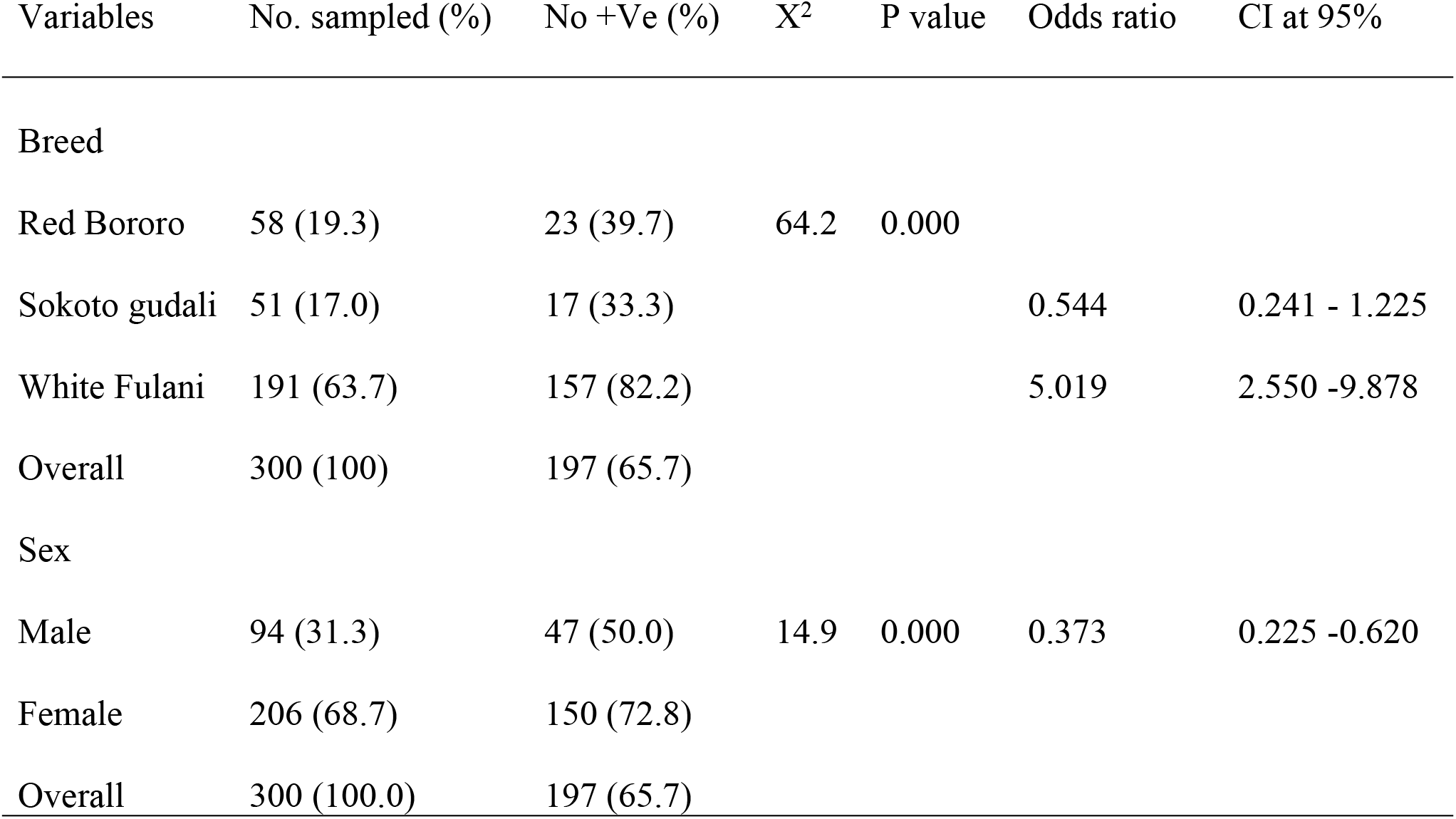
Seroprevalence of foot and mouth disease virus in cattle around Yankari game reserve and Sumu Wildlife Park in Bauchi State, Nigeria

Comparison of the overall seroprevalences of FMDV at the wildlife-cattle interface (Table 2) showed that detectable antibodies to FMDV were significantly higher (P < 0.05) in cattle 197 (65.67%) than in wildlife 13 (24%).

**Table 2:**
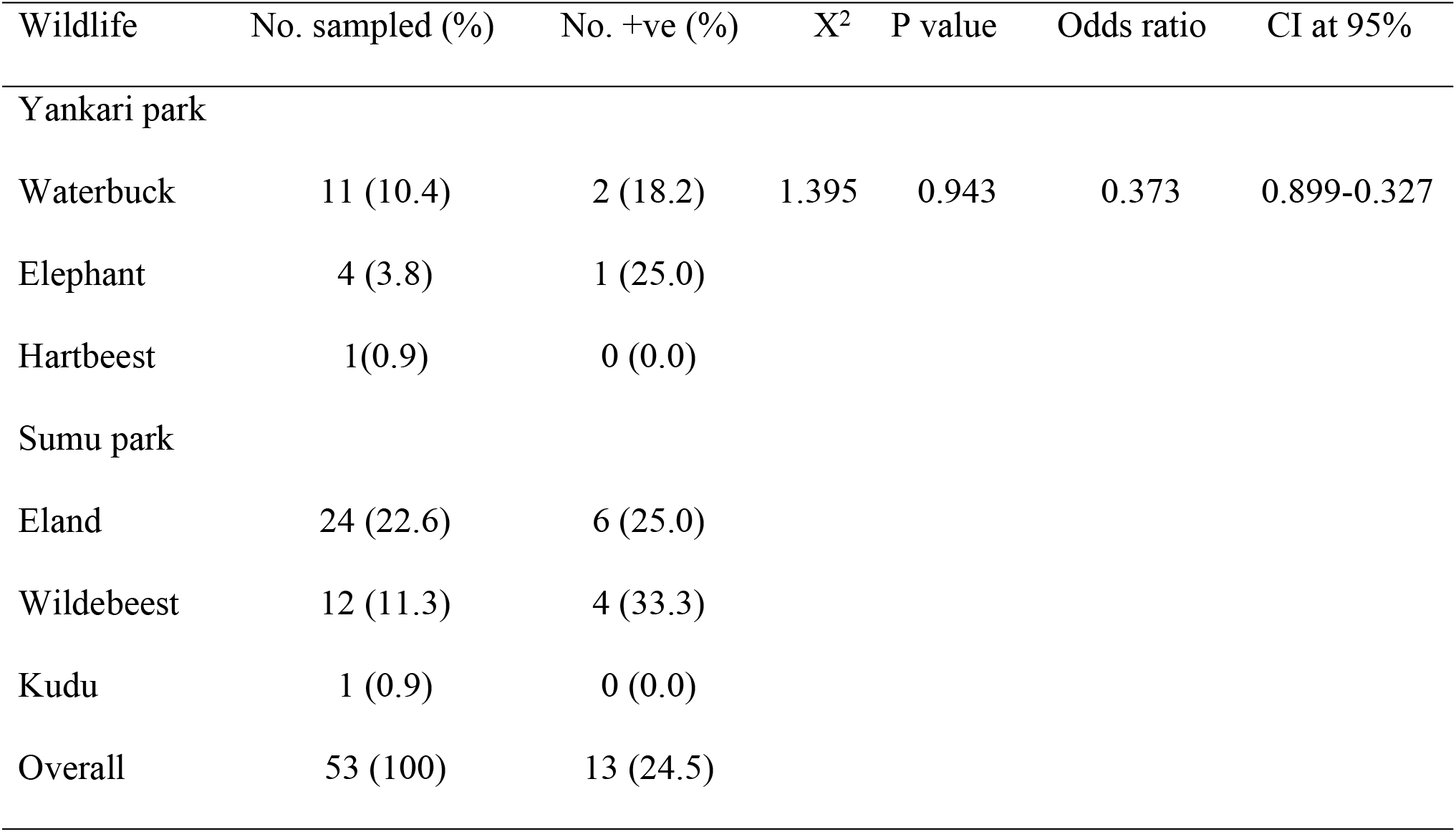
Seroprevalence of foot and mouth disease at the virus in wildlife from Yankari game reserve and Sumu wildlife park in Bauchi State, Nigeria.

Antibodies to FMDV were significantly higher in female cattle than males (P <0.05) with Bunaji breed of cattle having high risk factor (odds ratio >5) of exposure to FMDV than the other breeds of cattle examined (Table 3).

**Table 3:** Seroprevalence of foot and mouth disease at the wildlife-cattle interface in Yankari game reserve and Sumu wildlife park in Bauchi State, Nigeria.

| Specie | No. sampled | No. +Ve (%) | X<sup>2</sup> | P value | Odds ratio | CI at 95% |
| --- | --- | --- | --- | --- | --- | --- |
| Wildlife | 53 | 13 (24.53) | 31.63 | 0.000 | 0.1699 | 0.087 - 0.332 |
| Cattle | 300 | 197 (65.67) |  |  |  |  |
| Overall | 353 | 210 (59.49) |  |  |  |  |

The detectable antibodies to FMD serotype were for serotypes O, A, SAT 1 and SAT 2 in waterbuck, wildebeest and eland whereas antibodies to serotypes A and SAT 2 were detected in elephant. Each of the serotypes A and SAT 1 were shown to have highest reactors of 10 (18.87%) whereas serotype O had the least reactors of 7 (13.21%) (Table 4)

**Table 4:**
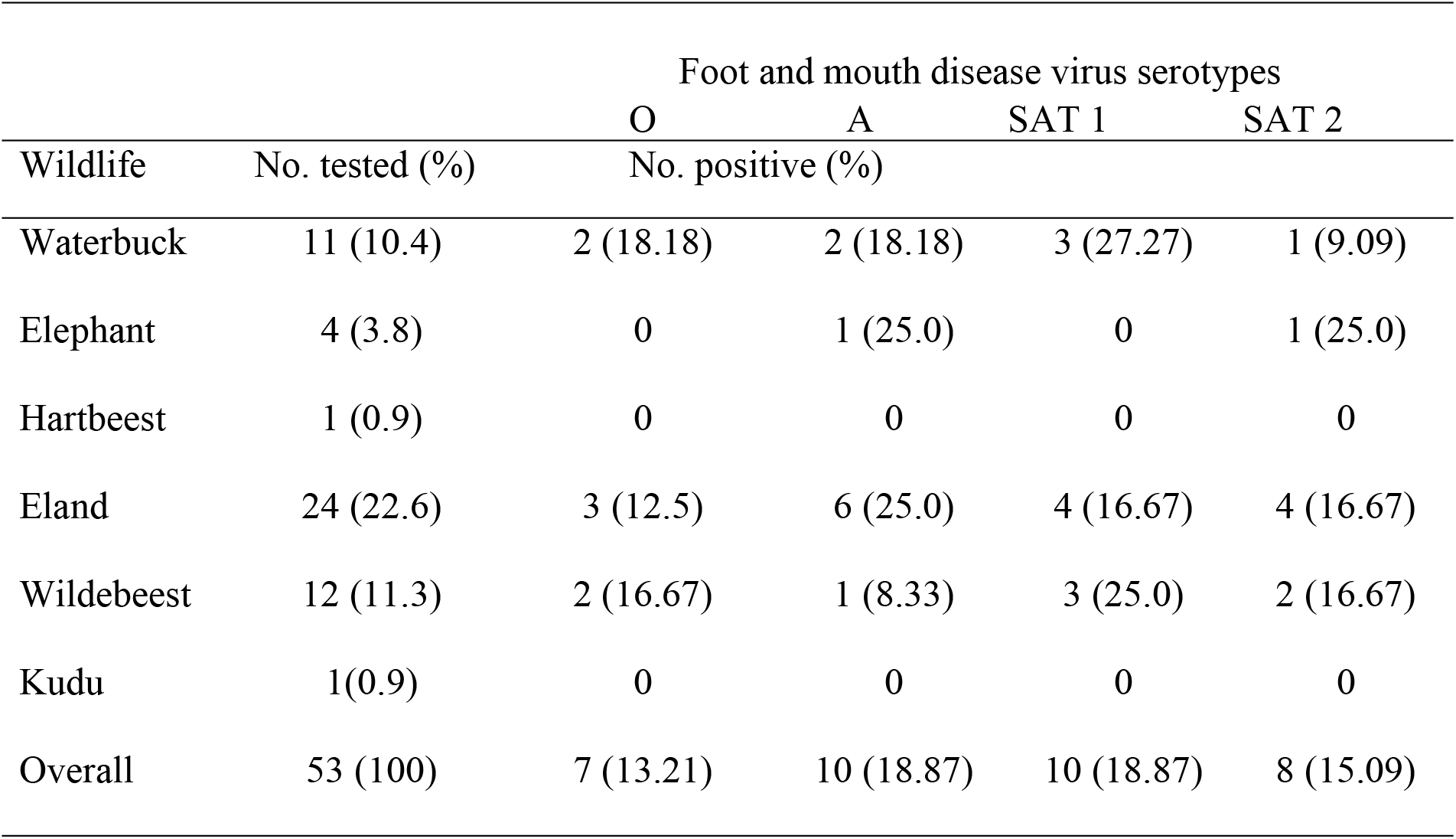
Foot and mouth disease virus serotypes detected in wildlife in Yankari game reserve and Sumu Wildlife Park in Bauchi State, Nigeria.

## Discussion

The results of this study have shown that antibodies to FMDV were present in cattle (65.7%) and wildlife (24.5%). This is consistent with results of previous survey for FMDV antibodies in Nigeria in which a seroprevalence of 75.11% was reported in a study conducted in cattle in Kwara State (34). Also, seroprevalences of 64.3% and 70.98% respectively were reported in studies carried out in Plateau State (35, 36), and 64.7% in a study conducted at the Border States in Nigeria (21, 37). The similarities of findings of the present study with previous studies have shown that FMD is still an enzootic disease in Nigeria and this could be attributed to the lack of FMD vaccination campaigns in Nigeria (21, 37). There is unrestricted herds mobility, continuous contact and intermingling of different cattle herds at water points, communal grazing areas and porous borders.

The higher FMDV seroprevalence in female cattle during this study was consistent with the findings of other investigators (34, 37) who reported a risk difference in association with sex during FMDV studies in Kwara and Plateau States, Nigeria, respectively. Similarly, high incidence of FMDV in females in Northwest Ethiopia was reported (38). However, most of the cattle sampled during the study were females as opposed to males. The significant association of seroprevalence with sex could be attributed to the preference for females to males by the nomads for reproductive purposes and milk production and therefore females are kept for longer period thereby having higher risk of exposure than males (8, 34, 37). Significant association in seropositivity was observed in Bunaji breed of cattle, this could be due to small number of other breeds (Sokoto gudali and Red bororo) sampled. However, all the breeds of cattle are equally at risk.

Results from the study have shown that antibodies to FMDV were present in elands, wildebeests, waterbucks and elephants. This finding being the first of its kind in the study area reveals that FMD could be a problem in wildlife in Nigeria. This is not surprising as FMD is endemic in Nigerian livestock (18, 20, 39, 23). Presence of wildlife population along the national park in Borgu Niger State Nigeria where cloven hoofed species come in contact with live stock was shown to be the probable exposure factor that contributed to high FMD sero-positivity in livestock observed in the area (37). The results from this study corroborate with other studies in South Africa, Zimbabwe, Zambia, Botswana, Namibia, India, Chad and Iran that demonstrated FMDV antibodies in wildlife (40, 10, 41, 42, 43, 28, 11, 44, 45). High FMDV prevalence in waterbuck observed in this study reflects their ecology and living ecosystem which is consistent with other findings in East Africa and Zimbabwe (41, 27, 46). The study hitherto provided a picture of FMDV distribution in wildlife in Bauchi State, Nigeria which was observed to be largely understudied (44).

The study confirms the presence of antibody to FMDV serotype O, A, SAT 1 and SAT 2 in wildlife a finding probably first of its kind in Nigeria. Reported outbreaks affecting livestock of West Africa since 2000 were caused by FMDV types O, A, and SAT 2 (44). Similarly, FMDV serotypes O, A, and SAT 2 were the cause of most reported outbreaks in domestic livestock in Nigeria from 2010 to 2016 (39, 34, 22). The result here showed that FMDV serotypes observed in wildlife were equally previously observed in domestic livestock. The possible source of FMDV serotypes infection for the wildlife could be from infected livestock interacting with wildlife in the same environment. Transmission of FMDV between wildlife and livestock, even in isolated areas, may be due to windborne infection or via fomites (47, 48). Wildlife species often congregate at the natural ‘salt lick’ point in YGR (31) similarly artificial salt lick points are also available in SWP. Therefore, dissemination of the FMDV during wildlife activities at the salt-lick points is possible. Previous studies have shown that FMDV can easily be disseminated in the soil and can persist in that environment for a long period (28).

The presence of FMDV antibodies in wildlife and cattle in this study might be driven by direct contact at wildlife-livestock interface through sharing of water and pasture resources observed to be a common activity in YGR and SWP in Bauchi State, Nigeria (31, 29). During dry season wildlife and livestock in the study area do closely congregate at feed and water points thus increasing the transmission likelihood of water-related infections like FMD (41, 13, 44). Studies conducted in Ethiopia and Zimbabwe found significant association between cattle exposed to FMDV and their contact history with wildlife (50, 48, 11). It is unfortunate that due to the endemic nature of FMD in Nigeria that outbreaks are not being investigated to determine the primary source and hence the disease have continued to be a scourge to live stock production in the Country.

## Conclusion

According to the results presented, presence of FMDV antibodies in cattle and some wildlife were observed with four serotypes of FMDV: O, A, SAT-1 and SAT-2 detected from the 3-ABC positive reactors in some wildlife. This might have been driven by direct contact at wildlife-cattle interface through sharing of water and pasture resources observed to be a common activity in YGR and SWP in Bauchi State, Nigeria. The study highlights the need for intensification of FMD surveillance system, control and prevention program among wildlife and livestock within and around wildlife conservation areas in Nigeria.

## Acknowledgements

We would like to thank the wildlife capture team and game rangers from Yankari Game Reserve and Sumu Wildlife Park in Bauchi State, Nigeria for their cooperation and support during samples collection.

